# Phosphorylation of the overlooked tyrosine 310 regulates the structure, aggregation, and microtubule- and lipid-binding properties of Tau

**DOI:** 10.1101/2020.01.06.896381

**Authors:** Nadine Ait-Bouziad, Anass Chiki, Galina Limorenko, Shifeng Xiao, David Eliezer, Hilal A. Lashuel

## Abstract

The microtubule-associated protein Tau is implicated in the pathogenesis of several neurodegenerative disorders, including Alzheimer’s disease. Increasing evidence suggests that post-translational modifications play critical roles in regulating Tau normal functions and its pathogenic properties in Tauopathies. Very little is known about how phosphorylation of tyrosine residues influences the structure, aggregation, and microtubule- and lipid-binding properties of Tau. In this work, we aimed to address this knowledge gap and determine the relative contribution of phosphorylation of one or several of the five tyrosine residues in Tau (Y18, Y29, Y197, Y310 and Y394) to the regulation of its biophysical, aggregation and functional properties. Towards this goal, we used a combination of site-specific mutagenesis and *in vitro* phosphorylation by c-Abl kinase to generate Tau species phosphorylated at all tyrosine residues, all tyrosine residues except Y310 or Y394 (pTau-Y310F, pTau-Y394F) and Tau phosphorylated only at Y310 or Y394 (4F\pY310 or 4F\pY394). Our results show that phosphorylation at all five tyrosine residues, multiple N-terminal tyrosine residues (Y18, Y29 and Y197) or site-specific phosphorylation at residue Y310, itself located in the microtubule-binding and aggregation-prone domain of Tau, was sufficient to abolish Tau aggregation and inhibit its microtubule- and lipid-binding properties. NMR studies demonstrated that these effects were mediated by a local decrease in β−sheet propensity of the PHF6 domain. Our findings underscore the unique role of Y310 phosphorylation in the regulation of Tau aggregation, microtubule and lipid interactions and highlight the importance of conducting further studies to elucidate its role in the regulation of Tau normal functions and its pathogenic properties.

## Introduction

The misfolding and aggregation of the microtubule (MT)-associated protein Tau have been linked to the pathogenesis of several neurodegenerative disorders (1), including Alzheimer disease (AD) (2), corticobasal degeneration (CBD) (3), progressive supranuclear palsy (PSP) (3,4), Pick’s disease (PiD), argyrophilic grain disease (AGD) (5), chronic traumatic encephalopathy (CTE) (6) and frontotemporal dementia with parkinsonism linked to chromosome 17 (FTDP-17) (7). In the brain of individuals affected by AD, aggregated and abnormally hyper-phosphorylated and modified forms of Tau, such as paired helical filaments (PHFs) and straight filaments (SFs), are found as the main component of the neurofibrillary tangles (NFTs), one of the key diagnostic pathological hallmarks of AD, in addition to amyloid plaques (8).

Tau is extensively post-translationally modified (PTM) with phosphorylation being one of the predominant modifications (9). Although many phosphorylation sites of Tau identified in NFTs are also found in healthy neurons, at advanced disease stages Tau aberrant hyper-phosphorylation (10) encompasses more than 39 potential sites (11,12), leading to a total phosphorylation level of Tau that is three to four times higher than that of normal Tau (13,14). The presence of hyper-phosphorylated Tau in both soluble cytosolic forms and highly aggregated and insoluble forms within NFTs (13,15) suggests that the pattern of hyper-phosphorylation or site-specific phosphorylation may play key roles in regulating its normal function(s) and/or pathogenic properties.

Serine and threonine residues greatly outnumber tyrosine residues in the Tau sequence, which explains why phosphorylation at these residues has received predominant attention. Tau contains five tyrosine residues (Fig. 1), located at positions Y18, Y29, Y197, Y310 and Y394, which are all susceptible to phosphorylation. Tau phosphorylation on tyrosine residues has been described only recently and remains understudied. A few tyrosine kinases have been shown to phosphorylate tyrosine residues in Tau both *in vitro*, in cell culture and *in vivo*. For example, Fyn (16,17) and Lck (17), members of the Src kinase family, phosphorylate Tau on all five tyrosine residues, however somewhat preferentially on residue Y18; while members of the c-Abl kinase family (c-Abl and Arg) were reported to phosphorylate Tau on Y394 (18,19) and to a lesser extent on Y197 and Y310. Other non-SH3-containing tyrosine kinases have also been reported to phosphorylate Tau on residue Y18, such as Syk (20) and Pyk2, the focal adhesion kinase (FAK) (21); and on residue Y197 by TTBK1 (22). Additionally, it was shown that an increase in Tau tyrosine phosphorylation correlated with the formation of Tau aggregates (23). Furthermore, several reports have established that Abelson tyrosine kinase (c-Abl) becomes active in a variety of neurodegenerative diseases, including Parkinson’s disease (PD) and several Tauopathies (24-26), suggesting a likely role in the regulation of Tau tyrosine phosphorylation levels in disease.

**Figure 1.**
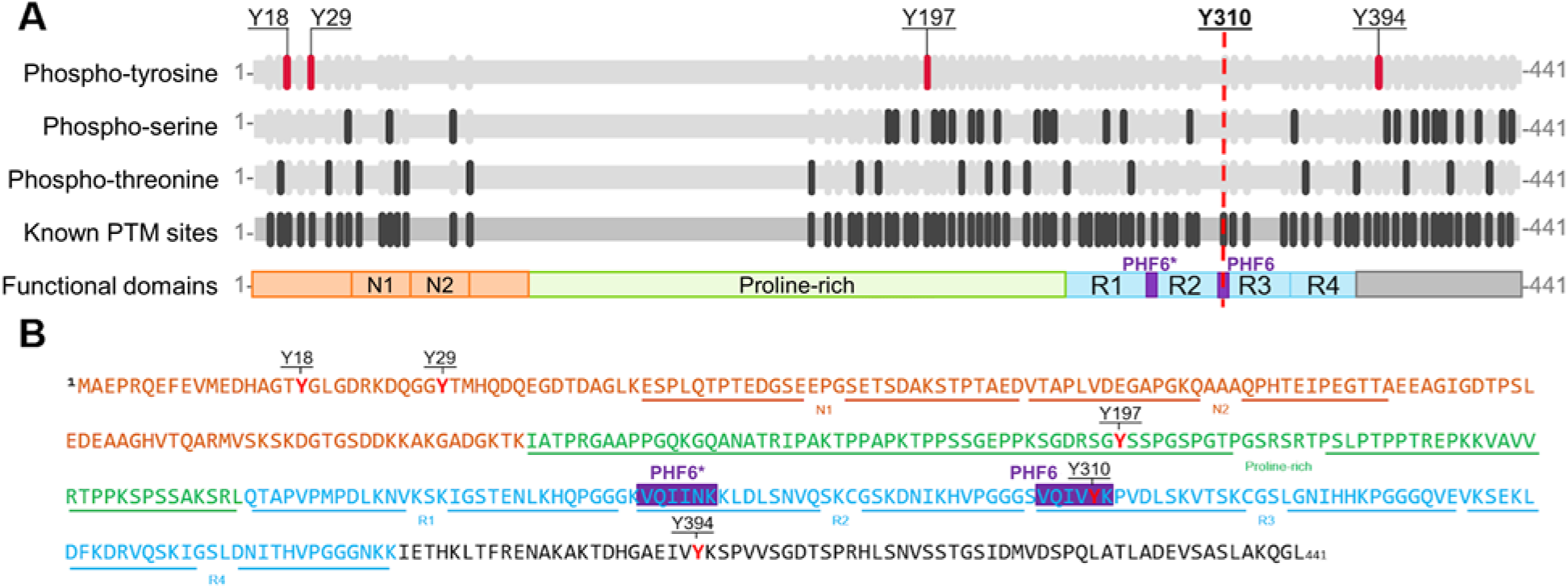
Schematic depiction of the sequence and different domains in Full-length Tau-441. **(A)** Map and domains of full-length Tau with known post-translational modification sites (grey); phospho-tyrosines (pY, magenta), phospho-serines (pS, dark grey) and phospho-threonines (pT, dark grey). Furthermore, Tau contains different functional domains: N-terminal sites (N1 aa 45-74, N2 aa 75-103), a proline-rich region (aa 151-224), and four repeat domains (R1 aa 244-274; R2 aa 275-305; R3 aa 306-336; R4 aa 337-368). Microtubule-binding region (MTBR) contains 2 hexapeptide domains PHF6* (aa 275-280, purple) and PHF6 (aa 306-311, purple). Adapted from http://www.Tauptm.org (78). **(B)** The annotated sequence of full-length Tau. Tau possesses 5 tyrosine residues, as indicated in red, at positions Y18, Y29, Y197, Y310 and Y394. Y18 and Y29 are localized in the N-terminal region of Tau (orange, N-terminal repeats N1 and N2 underlined), Y197 in the proline-rich domain (green), Y310 in the third repeat of the MTBR of Tau (blue) and Y394 in the C-terminal part of the protein (black). PHF6 and PHF6* hexapeptides are indicated in purple.

Alternative splicing of exon 2, 3 and 10 exons of Tau produces 6 isoforms from chromosome 17, the splicing results in an assembly domain that contains three or four microtubule-binding repeats (27). The four repeats form the proteolysis-resistant core of Tau aggregates (28). PHF6* and PHF6 hexapeptides (Fig. 1, A and B) are important elements for Tau binding to microtubules and constitute the minimal motif allowing fibrillization (29,30). K19 and K18 are fragments of Tau that comprise three and four repeat domains, respectively. These fragments have high aggregation and microtubules-binding propensities (31,32). Whereas phosphorylation of serine and threonine residues exhibit a degree of clustering in the N- and C-terminal functional domains encompassing the repeat regions and the MT-binding domains 1 and 2 (Fig. 1, A and B), there is only one tyrosine phosphorylation site (Y310) within the MT-binding domain. This residue is located in the repeat peptide R3, which has also been shown to play key roles in regulating Tau aggregation, PHF formation, and membrane interactions (33,34).

Interestingly, despite the prominent position of Y310 within the aggregation-prone hexapeptide PHF6 (amino acids 306-311), only a handful of studies have specifically investigated the roles of this residue in regulating Tau structure, aggregation, and toxicity. Importantly, none of the previous studies (30,35-42) specifically investigated the effects of Y310 phosphorylation in the context of the full-length Tau protein, relying instead on the short PHF6-containing peptides, featuring PHF6-derived sequences as short as 3 amino acids (43). Santa-Maria *et al*. (44) showed that phosphorylation of Y310 abolished the aggregation of the PHF6 fragment (amino acids 306-311) *in vitro*. Conversely, Inoue *et al*. reported contradictory results, where phosphorylation of Y310 resulted in a marked increase in the propensity of PHF6 peptide to form fibrils (40,45,46). Furthermore, while previous studies have suggested that Tau phosphorylation on tyrosine residues is an important modulator of Tau functions under both normal conditions and in the course of AD pathogenesis (16,47), the role of each tyrosine residue in regulating Tau protein aggregation, MT-binding, and lipid-binding propensities has not been systematically investigated.

This knowledge gap combined with the location of Y310 within the PHF6 domain of Tau prompted us to conduct more detailed investigations to 1) determine the effects of tyrosine phosphorylation on the regulation of the normal function and pathogenic properties of full-length Tau; 2) elucidate the relative contribution of phosphorylation at each tyrosine residue; and 3) investigate the effect of phosphorylation at Y310 on the structure, aggregation and microtubule-binding of the full-length Tau and microtubule-binding domain-containing Tau K18 fragment. Towards achieving these goals, we utilized a combination of mutagenesis and *in vitro* phosphorylation using the c-Abl kinase to produce milligram quantities of Tau phosphorylated at single or multiple tyrosine residues. Our findings show that phosphorylation of Y310 is sufficient to inhibit Tau aggregation and plays a key role in regulating Tau microtubule-binding properties and interactions with the membrane, thus underscoring the importance of further studies to elucidate the role of phosphorylation at this residue in regulating Tau function under normal and pathological conditions.

## Results

### c-Abl phosphorylates Tau on multiple tyrosine residues *in vitro*

To investigate the role of tyrosine phosphorylation in regulating Tau structure, aggregation and MT-binding properties, we first sought to develop conditions that allow for efficient phosphorylation of Tau *in vitro*. Towards this goal, we assessed the phosphorylation efficiency and specificity of c-Abl kinase for Tau. Recombinant full-length Tau was subjected to *in vitro* phosphorylation using recombinant c−Abl and the extent of phosphorylation and number of phosphorylation sites was systematically followed by LC/MS. Fig. 2, A shows that c-Abl phosphorylated Tau at multiple sites, up to 5 sites after 4 h of incubation in reaction buffer. Increasing the amount of kinase or incubation time did not cause a shift towards species with a higher number of phosphorylated tyrosine residues or relative phosphorylation levels (data not shown). Although RP−HPLC allowed the separation of the kinase from the phosphorylated Tau proteins, it was not possible to achieve separation of the different phosphorylated Tau species, which eluted in one peak as observed by the UPLC and a single band by SDS-PAGE (Fig. 2, B and C). To determine the exact tyrosine phosphorylation profile of Tau, the purified mixture of pTau was digested by trypsin and the resulting peptides were analyzed by LC/MS/MS. The analysis of the phosphorylated sites was performed using the Scaffold software (http://www.proteomesoftware.com/) which determines the abundance of phosphorylated peptides based on the spectral counting (SP). As shown in Fig. 2, D and E, c−Abl phosphorylated Tau on Y18, Y29, Y197, Y310 and Y394, with Y310 and Y394 being the major phosphorylation sites.

**Figure 2.**
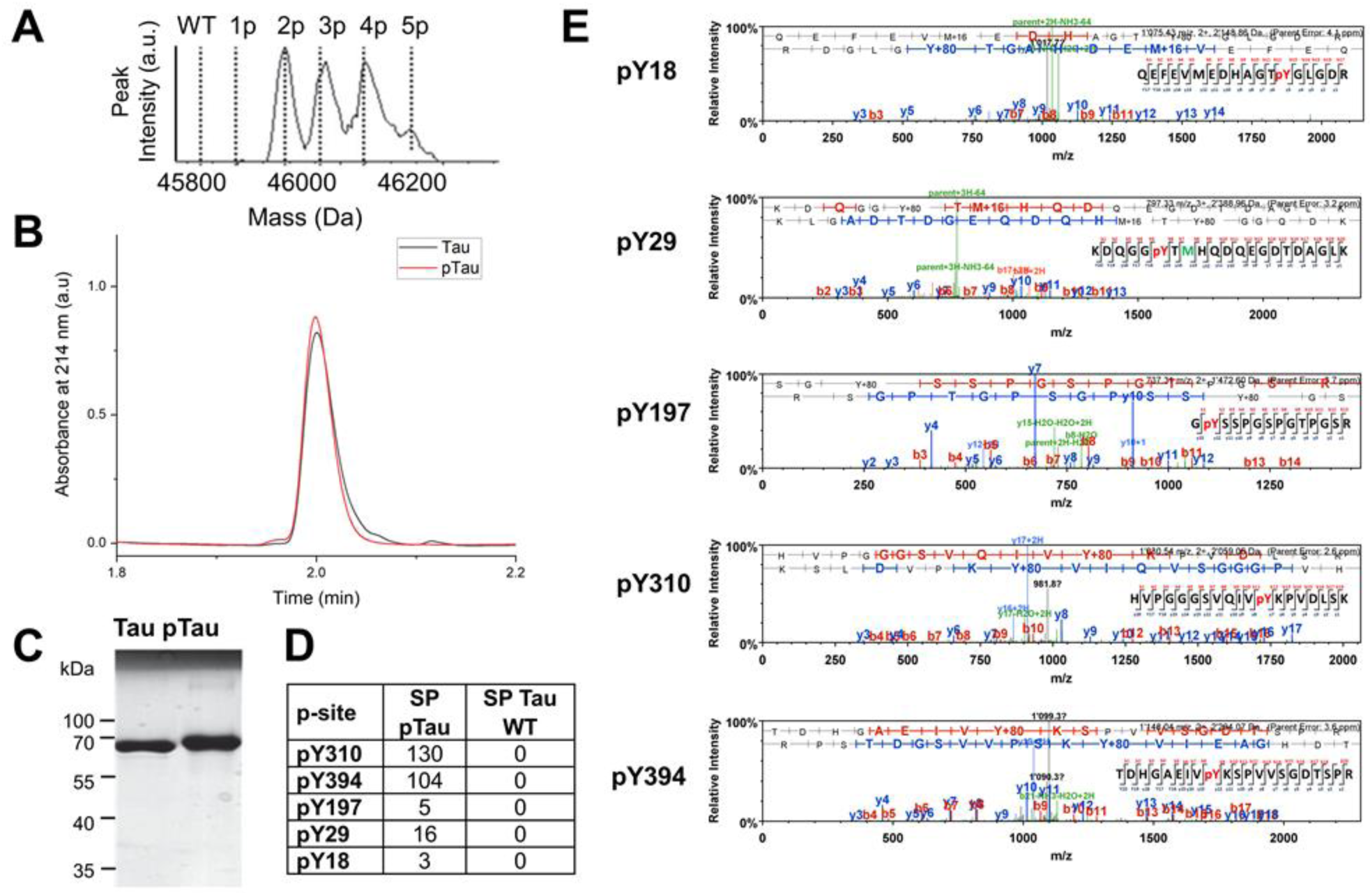
c-Abl-mediated *in vitro* phosphorylation and characterization of Tau. (pTau); phosphorylation was performed for 4 h, which leads to a mixture of Tau phosphorylated on two to five residues. Characterization of the RP-HPLC purified Tau and the pTau mixtures by LC-MS **(A)**, UPLC **(B)** and SDS-PAGE **(C). (D and E)** Tandem LC/MS/MS of RP-HPLC purified pTau following trypsin digestion and peptides enrichment using a TiO_2_ resin. The analysis of the phosphopeptides was performed using the Scaffold version 4.0 (Proteome Software), and the number of phosphopeptides detected per phosphorylation site was reported as the spectral count (SP).

### c-Abl phosphorylation of Tau significantly delays its aggregation

To determine the effect of tyrosine phosphorylation on Tau aggregation, we carried out *in vitro* phosphorylation of Tau using the c-Abl kinase and purified the mixture of Tau phosphorylated proteins using RP-HPLC. Next, we compared *in vitro* aggregation of the non-phosphorylated to c-Abl-phosphorylated Tau (Fig. 3) by incubating and monitored the aggregation of both proteins (at10 µM) over two days at 37°C under shaking conditions, in presence of 2.5 µM heparin. We found that phosphorylated Tau (pTau) was unable to form canonical β−sheet-containing fibrils, as demonstrated by 1) electron microscopy (Fig. 3A); 2) the absence of conversion of Tau conformation to β−sheet structure as discerned by circular dichroism (CD) (Fig. 3, B); and 3) the fact that the ThT fluorescence remained at baseline levels (Fig. 3, C). However, large oligomers were discernible by electron microscopy at 12 h, and longer fibrils were apparent only after 24-48 h incubation. These observations were further confirmed by the fact that the sedimentation assay showed an increase of the SDS-resistant oligomeric species overtime in the pTau sample, but not for the unmodified protein (Fig. 3, A, D).

**Figure 3.**
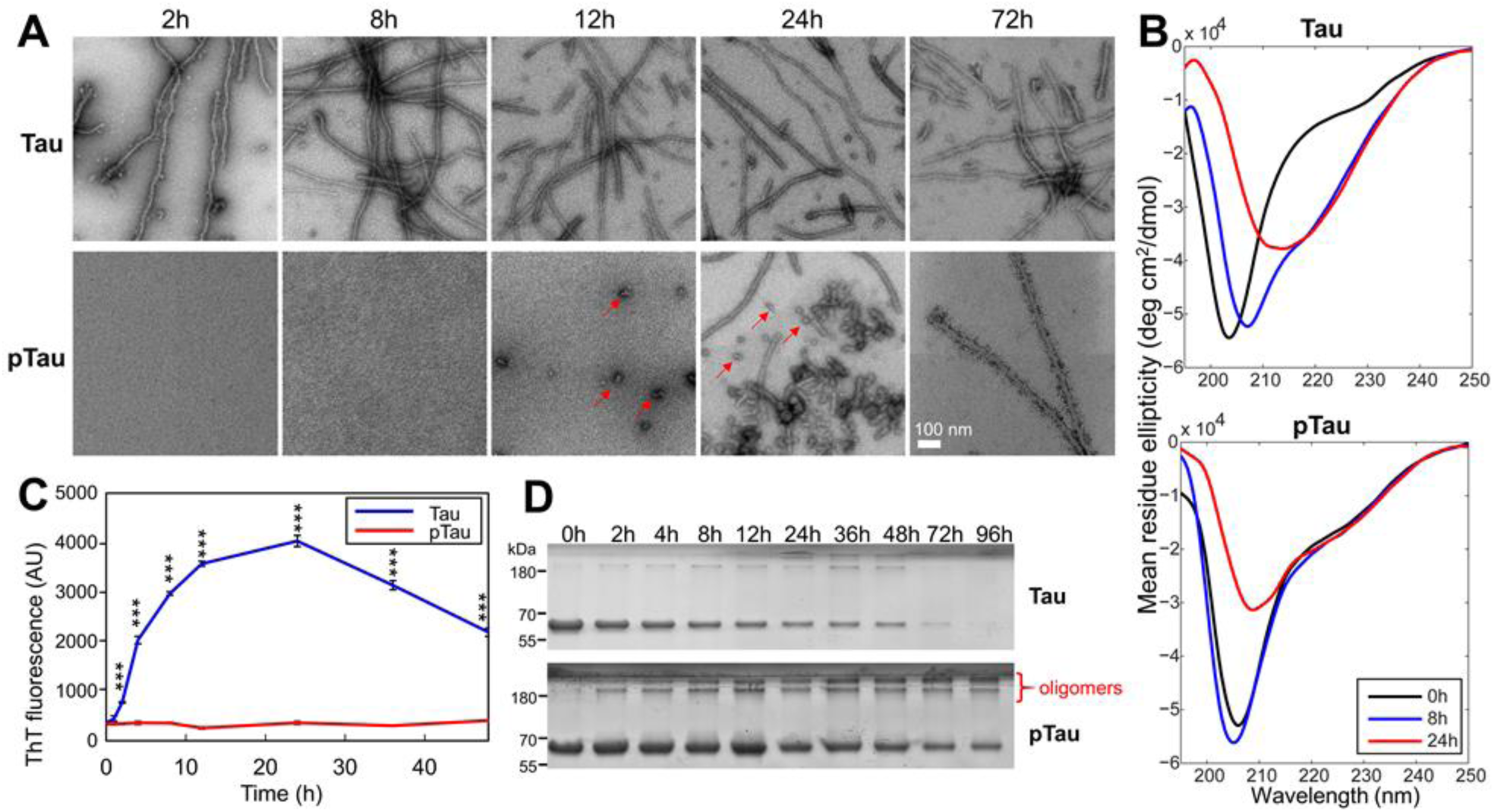
Comparison of the aggregation properties of WT Tau and purified c-Abl phosphorylated Tau, pTau. 10 µM Tau and pTau were incubated for 48 hours at 37°C under shaking conditions, in presence of 2.5 µM heparin and the extent of aggregation was monitored by EM (scale bar = 100 nm for all images) **(A)**, CD spectroscopy **(B)** ThT fluorescence **(C)**, and sedimentation assays **(D)**. Quantification of the supernatant band intensity could not be performed due to the presence of soluble non−pelletable oligomeric species in the pTau sample. In **(C)** Two-tailed ANOVA with Sidak’s multiple comparisons test, significance values are indicated by *** p<< 0.001; time points at 0h and 1h were not significantly different.

It is likely that oligomeric species observed in the unmodified Tau sample were ON-pathway to fibril formation and represent intermediate species that disappeared by the end of the reaction through equilibrium shift towards fibrillar structures. In contrast, the oligomers formed by pTau might represent OFF-aggregation-pathway species that did not proceed to form fibrils and remained in the sample at 96h (Fig 3, D). These results demonstrate that phosphorylation of Tau at multiple tyrosine residues significantly delays its aggregation and fibril formation and results in the accumulation of phosphorylated Tau oligomers.

### c-Abl-mediated inhibitory effects on Tau aggregation depend on the site of tyrosine phosphorylation

To determine the relative contribution of each tyrosine residue to the c-Abl phosphorylation-mediated inhibition of Tau aggregation, we generated Tau proteins that were phosphorylated or not at one or a selected number of tyrosine residues focusing on Y394 and Y310. To allow site-specific phosphorylation of these residues, we mutated all tyrosine residues except Y310 (4F\Y310) or Y394 (4F\Y394) to phenylalanine. In addition, we generated Y310 and Y394 residue-specific phosphorylation deficient Tau mutants, where each of these residues is mutated to phenylalanine to allow phosphorylation of all other tyrosine residues except at Y310 (Y310F) or Y394 (Y394F) (Fig. 4, A, hollow circles). These Tau variants were subjected to *in vitro* phosphorylation using c-Abl (Fig. 4, A, red circles) and the phosphorylated proteins were purified and their aggregation properties were compared to that of the corresponding non-phosphorylated variants. For the 4F\Y310 and 4F\Y394 mutants, which could only be phosphorylated at Y310 and Y394, complete phosphorylation was achieved, whereas a mixture of up to four phosphorylation sites was observed for the single Y310F and Y394F mutants (see LC/MS spectra in Table 1). We next investigated the kinetics of aggregation of Tau Y−>F mutants using the same experimental conditions as for full-length Tau (Fig. 4). We observed that 4F\pY310, pTau−Y310F and pTau−Y394F were unable to aggregate, as demonstrated by the lack of increase in the ThT fluorescence for these samples (Fig. 4, B), while 4F\pY394 retained its capacity to form β−sheet-containing species. These findings were supported by EM data (Fig. 4, D), where only the 4F\pY394 sample was found to contain Tau of fibrillar morphology, whereas all other phosphorylated mutants only showed the presence of oligomeric species (Fig. 4, D, arrows). These results were further supported by the sedimentation assay, whereupon removal of fibrillar species by ultracentrifugation monomeric Tau species showed time-dependent decrease in soluble Tau in all non-phosphorylated mutants (Fig. 4, C) that paralleled the increase in ThT fluorescence signal (Fig. 4, B), signifying the recruitment of monomeric Tau into β−sheet-containing species. A time-dependent decrease in monomeric Tau over time was also observed for phosphorylated Tau mutants. However, in contrast to their non-phosphorylated counterparts or mutant 4F\pY394, higher molecular weight Tau bands were apparent for phosphorylated Tau mutants 4F\pY310, pTau−Y310F and pTau−Y394F (Fig. 4, C, red boxes), suggesting the presence of Tau oligomeric species, which agrees with EM and ThT data. These findings suggest that phosphorylation at Y310, but not Y394 is sufficient to inhibit the fibrillization of Tau, as 4F\pY394 was able to aggregate to the same extent as its non−phosphorylated version. Interestingly, phosphorylation of the Y310F mutant also suppressed Tau fibril formation, even though in this mutant the residue Y310 was not available for phosphorylation, indicating that phosphorylation of multiple N-terminal tyrosine residues (Y18, Y29 and Y197) may also play important role in inhibiting Tau aggregation These results show that although Y310 phosphorylation is sufficient to suppress the aggregation of the full-length Tau, the interplay between phosphorylation of the other tyrosine residues is also of critical importance in modulating Tau oligomerization and fibril formation.

**Table 1.**
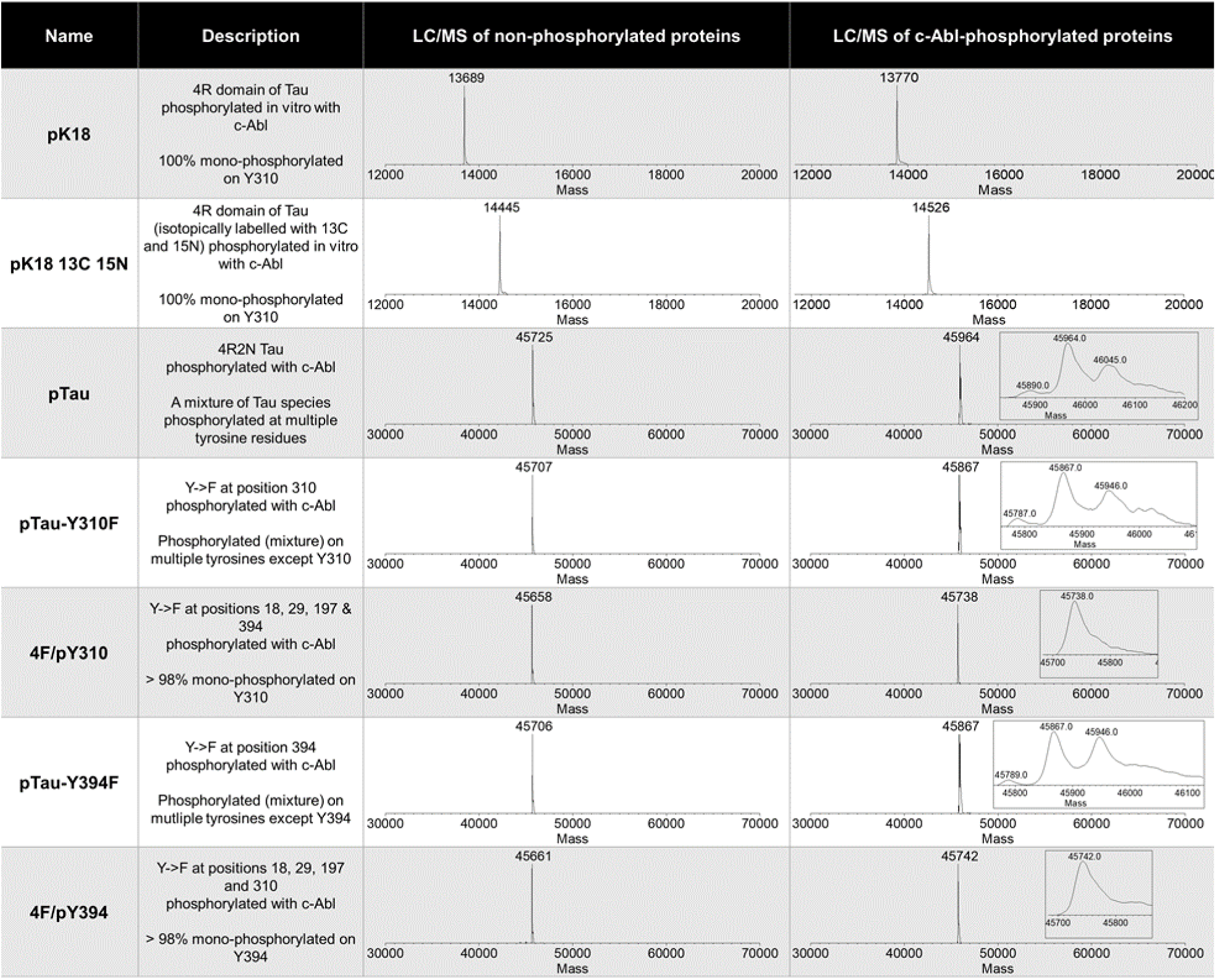
A list of the phosphorylated proteins used in this study, including how they were prepared and the LC/MS data for each protein before and after *in vitro* phosphorylation with c-Abl.

**Figure 4.**
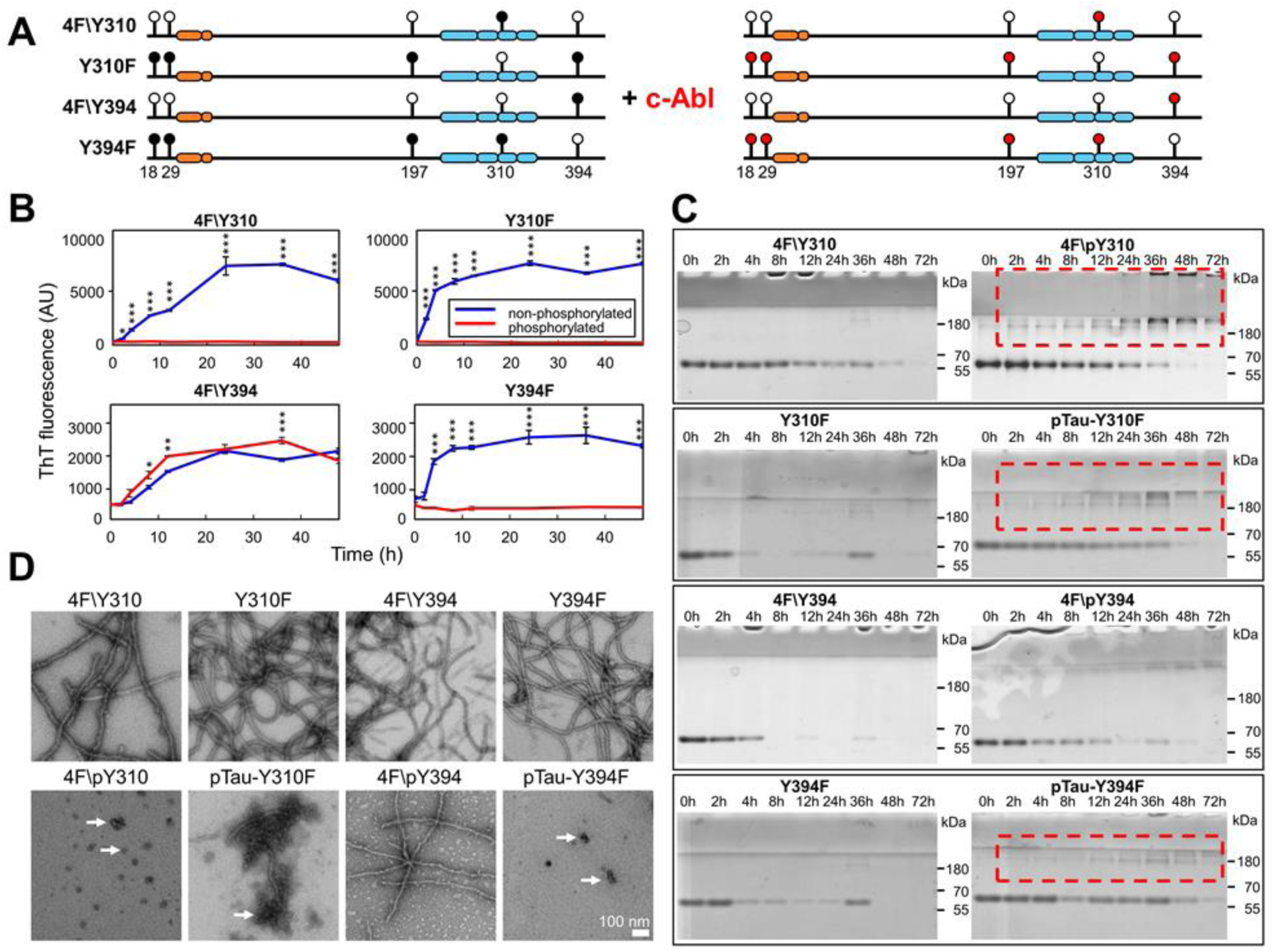
Comparison of the aggregation behavior of non-phosphorylated and phosphorylated Y->F Tau mutants. **(A)** Schematic depiction showing the position of Tau tyrosine residue Y−> F mutations (filled circle: Y; hollow circle: F) and the tyrosine sites phosphorylated by c-Abl (red circles). For the aggregation studies, 10 µM non- and phosphorylated Y−>F Tau mutants were incubated for 48 hours at 37°C under shaking conditions, in the presence of 2.5 µM heparin and the extent of aggregation was monitored by ThT fluorescence **(B)**, EM (scale bar = 100 nm for all images) **(C)**, and sedimentation assays **(D). (B)** Kinetics of aggregation of the Y−>F Tau mutants monitored by changes in ThT fluorescence at different time points (graphs show one representative experiment for each mutant). Of the phosphorylated proteins only the 4F\pY394 was able to form fibrils, as detected by the increase in ThT fluorescence over time. Two-tailed ANOVA with Sidak’s multiple comparisons test, significance values are indicated by * p<0.05, ** p< 0.01 and *** p<< 0.001. **(C)** At several time points, an aliquot of the aggregation was taken and centrifuged. The supernatants were run on SDS−PAGE. All of the non−phosphorylated mutants showed a significant reduction of the soluble fraction over time. Out of the phosphorylated proteins, only the 4F\pY394 presented a similar reduction in the amount of soluble protein over time, while the 4F\pY310, pTau−Y310F and pTau−Y394F formed non−pelletable oligomeric structures (red dashed rectangles). **(D)** EM micrographs of non-phosphorylated (top panels) and phosphorylated (bottom panels) Tau Y−>F mutants at 24 h. Of the phosphorylated proteins only the 4F\pY394 was able to form fibrils, while 4F\pY310, pTau−Y310F and pTau−Y394F formed many oligomeric and amorphous structures (arrows).

### Tyrosine 310 phosphorylation underlies the structural basis of the inhibitory effect on Tau aggregation

The strong inhibitory effects of Y310 phosphorylation combined with the fact that it represents the only tyrosine residue within the MT-binding domain of full-length Tau and K18 fragment prompted us to investigate the molecular and structural bases by which Y310 phosphorylation inhibits Tau aggregation propensity. Towards this goal, we performed nuclear magnetic resonance (NMR) studies on the K18 peptide, which encompasses the MT-binding region (residues 244-368 of Tau-441 sequence) and includes the highly amyloidogenic PHF6 and PHF6* hexapeptides, the former of which contains Y310 (Fig. 1). K18 is also known for its high propensity to aggregate *in vitro* and in cells (48).

To this end, we produced recombinant 13C 15N-labelled K18 and subjected it to *in vitro* phosphorylation using recombinant c−Abl under the same experimental conditions as for full-length Tau. Given that there is only one tyrosine residue in K18, we were able to achieve complete K18 phosphorylation on Y310 within 4 h and obtained highly pure Y310-specifically modified pK18 (Fig. 5, A-C). Next, to determine the effect of tyrosine 310 phosphorylation on the structural properties of K18, we performed NMR analysis of 15N-labeled pK18. The proton−nitrogen HSQC spectrum of pK18 compared to that of K18 showed that phosphorylation at residue Y310 induced local effects on the structure of the peptide, as indicated by chemical shift changes for signals originating from residues surrounding Y310 (Fig. 5, D). Interestingly, signals that were affected in the spectrum of pK18 consistently moved towards lower 15N chemical shifts. Because increases/decreases in NMR 15N chemical shifts correlate with increases/decreases in local β-strand structure (49), the observed decrease in 15N chemical shifts upon Y310 phosphorylation indicated a local decrease in β-sheet propensity for the PHF6 region as well as the subsequent ∼10 residues (Fig. 5, E). This effect likely results from the added negative charges of the phosphate group. Notably, the affected region adopts β-sheet conformations in all known Tau fibril structures. Thus, the local decrease in β-sheet propensity may have important implications in the context of Tau fibrillization, especially as the PHF6 hexapeptide exhibits the strongest intrinsic β-sheet propensity in the Tau sequence (50) and is critical for the aggregation of the protein (30,51).

**Figure 5.**
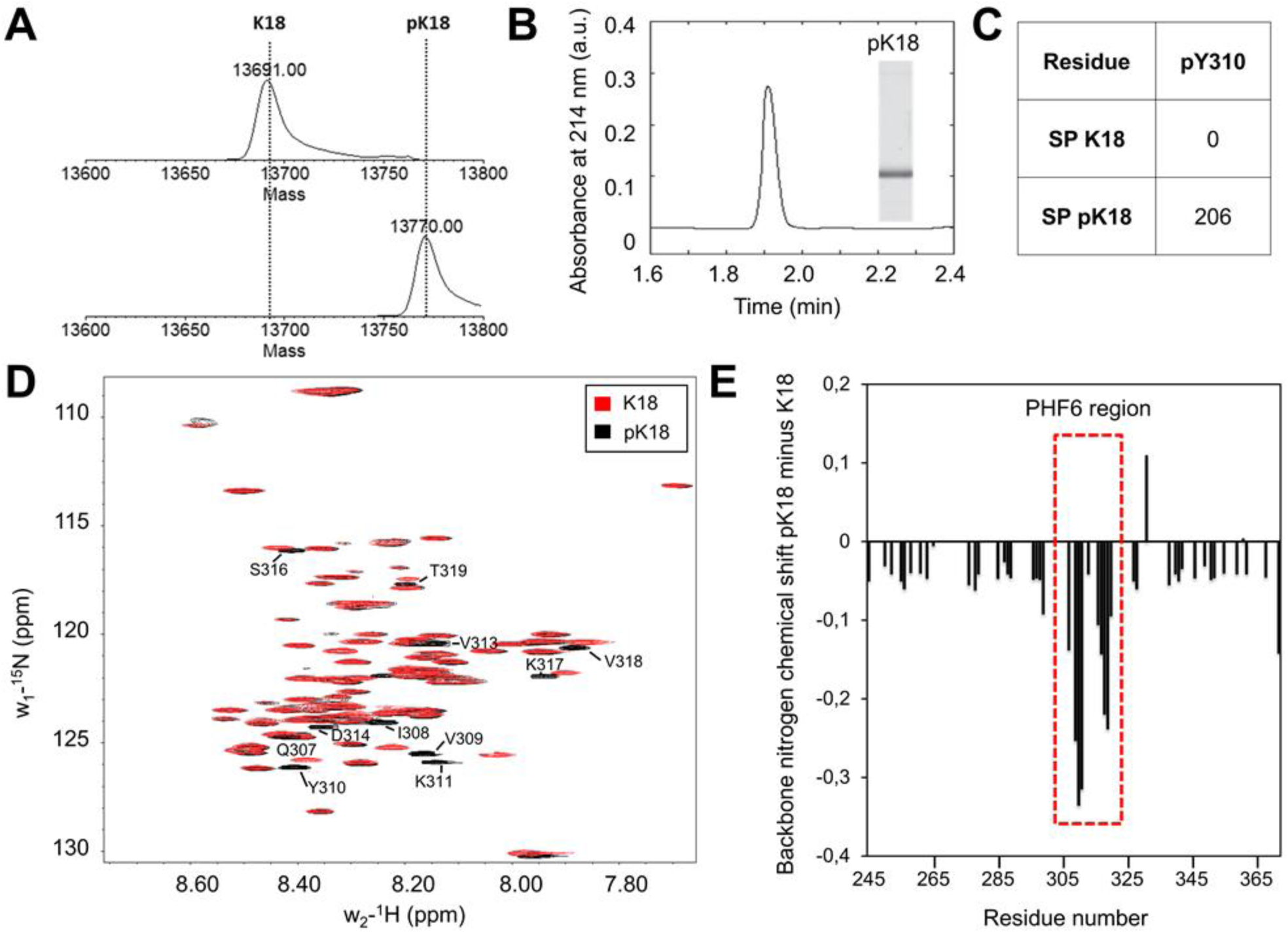
Detailed characterization of phosphorylated K18 at Y310 (pK18) and NMR analysis of 15N pK18 compared to the non-phosphorylated K18. LC/MS spectra **(A)**, UPLC profile and SDS-PAGE **(B)** of reverse-phase HPLC of purified pK18 following phosphorylation by c−Abl (phosphorylation was performed for 4 h). K18 phosphorylated by c−Abl leads to the complete phosphorylation at residue Y310. **(C)** The analysis of the phosphopeptides was performed using the Scaffold version 4.0 (Proteome Software), and the number of phosphopeptides detected per phosphorylation site was reported as the spectral count (SP). **(D)** Proton−nitrogen HSQC spectrum of unmodified K18 (red) compared to that of pK18 (black). Phosphorylation on Y310 induced local effects on the structure of pK18, as shown by chemical shift changes to annotated signals. **(E)** Decreased values of nitrogen chemical shifts (plotted as pK18 minus K18) indicate that phosphorylation decreases the β−sheet propensity of K18 in the PHF6 region, as well as for the subsequent ∼10 residues, all of which are found in a β-sheet conformation in Tau fibrils.

To further test whether Y310 phosphorylation impacted K18 aggregation, we compared the *in vitro* aggregation of Y310 phosphorylated K18 to that of WT K18(Fig. 6). As shown in Fig. 6 A and B, the aggregation of pK18 was significantly delayed compared to unmodified K18. In fact, both the ThT (Fig. 6, A) and the sedimentation assays (Fig. 6, D) showed a delay of about 2 h in the fluorescence increase and loss of soluble protein, respectively. Interestingly, the final absolute value of the plateau ThT signal of the phosphorylated K18 was consistently about two to three times higher than that of WT K18, suggesting a difference in fibril structure, packing of the fibrils or susceptibility to fragmentation (Fig. 6,C), which may affect ThT binding capacity and thus the end-point ThT value.

**Figure 6.**
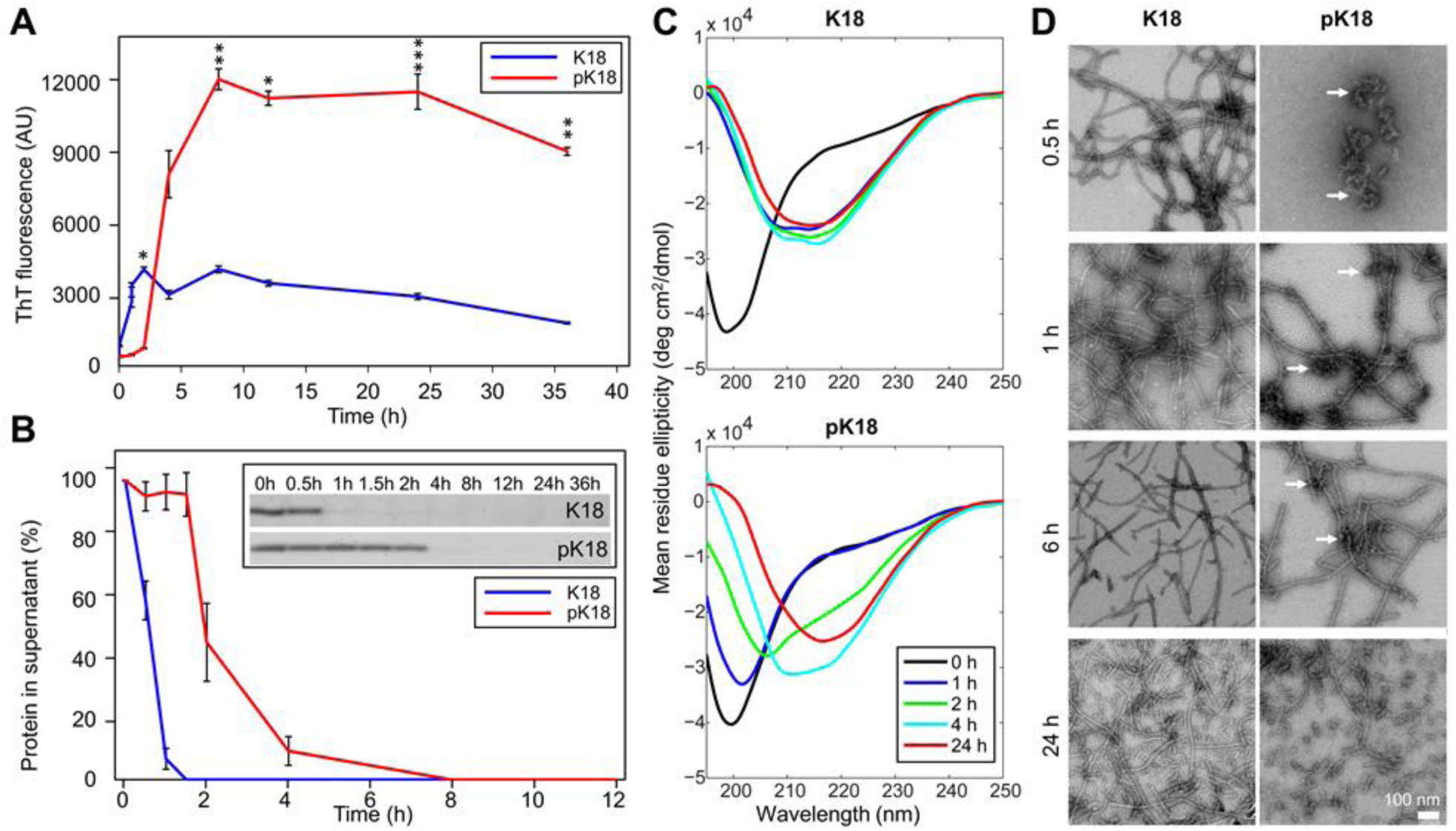
Comparison of the aggregation behavior of non-phosphorylated and phosphorylated K18. 10 µM non- and phosphorylated Y−>F Tau mutants were incubated for 48 hours at 37°C under shaking conditions, in the presence of 2.5 µM heparin and the extent of aggregation was monitored by ThT fluorescence **(A)**, sedimentation assays **(B)**, circular dichroism **(C)** and EM (scale bar = 100 nm for all images) **(D)**. Together, the results from these different assay show that phosphorylation at Y310 plays an inhibitory role during the nucleation phase of K18 aggregation, and delays formation of β−sheet-rich K18 fibrils. In **(A)** Two-tailed ANOVA with Sidak’s multiple comparisons test, significance values are indicated by * p < 0.05, ** p < 0.01, *** p<< 0.001. **(B)** At several times points, an aliquot of the aggregation reaction was taken and centrifuged. The supernatants were run on an SDS−PAGE (inlet). The supernatant band intensity from four independent experiments was quantified and reported as the percent of protein remaining in the supernatant (mean +/-SD).

To further investigate the effect of phosphorylation at Y310 on the early formation of β-sheet-containing species of K18, we monitored changes in K18 secondary structure overtime under aggregation conditions by CD spectroscopy. K18 displayed significantly higher ThT fluorescence signal than pK18 (Fig. 6, A) and sedimented more rapidly than pK18 (Fig 6, B). In addition, K18 fully converted to β−sheet within 1 h, whereas pK18 clearly showed markedly delayed conformational shift toward β−sheet spectra during 4 h of incubation (Fig. 6, C). As shown in Fig. 6 D, EM studies confirmed that fibril formation by pK18 was also delayed and showed significant accumulation of oligomeric amorphous aggregates at early and intermediate time points (Fig. 6, D, 0.5 h, 1 h, 6 h: arrows), and accumulation of fragmented and fully mature fibrillar morphologies, in addition to some oligomers at later time points (Fig. 6, D, 6 and 24 h). These findings suggest that Y310 phosphorylation results in delayed formation of β−sheet-rich K18 fibrillar structures (Fig. 6, A, B) and plays an inhibitory role during the nucleation phase of K18 aggregation, as demonstrated by attenuation of the pK18 conformational change (Fig. 6, C).

### Phosphorylation on tyrosine residues reduces Tau affinity for MTs

In light of the prominent direct association of Tau with MTs, we next asked whether Tau tyrosine phosphorylation, especially on residues Y310 and Y394, influence Tau microtubule-binding propensity. To address this question, we used full-length Tau phosphorylated by c−Abl, taking advantage of the preferential phosphorylation of Y310 and Y394 among other residues, as demonstrated by mass spectrometry SP counts (Figure 2, D). We first assessed the binding propensity of pTau to the MTs by quantifying the percent of protein pelleted when incubated with pre−formed paclitaxel−stabilized MTs at a single concentration (250 µg/ml Tau and 100 µg/ml of tubulin) (Fig. 7, A). We observed, as previously reported (16), that Tau bound to tubulin with about 50% efficiency, while this efficiency was decreased to approximately 30% in the case of pTau. As a negative control, the BSA was found to bind MTs with low efficiency of 12%, and as a positive control, the Microtubule−Associated Protein Fraction (MAPF) showed a high binding of 78% (Fig. 7, A), as expected. These results were further confirmed by the SDS-PAGE, where band corresponding to the molecular weight of non-phosphorylated Tau was enriched in the pellet fraction (Fig. 7, C, solid red arrows), whereas pTau remained predominantly in the soluble fraction. EM data showed no morphological differences for MTs bound to Tau or pTau (Fig. 7, B).

**Figure 7.**
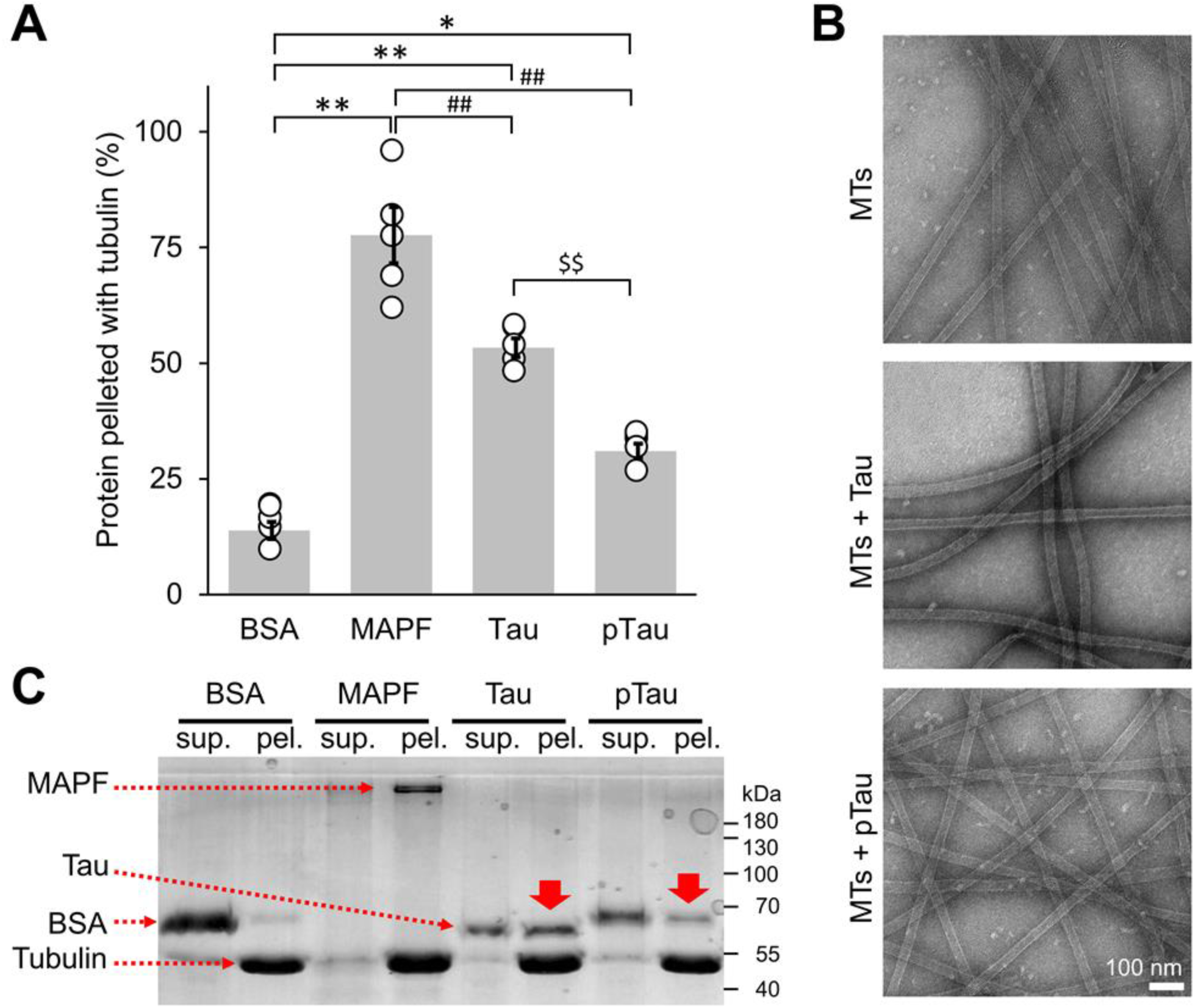
Tau/microtubule binding assay. **(A)** The binding propensity of pTau to the MTs was quantified as the percent of protein pelleted when incubated with pre−formed paclitaxel−stabilized MTs at a single concentration (250 µg/ml pTau and 100 µg/ml of tubulin). Tau bound to tubulin with about 50 % efficiency, while this efficiency was decreased to about 30% in the case of pTau. As a negative control, we used BSA, which showed low MT binding efficiency of 12%. As a positive control, the Microtubule−Associated Protein Fraction (MAPF) was used and showed a high binding of 78%. The averaged quantification from 5 repeats, represented as mean +/- SD with individual points plotted. Two-tailed ANOVA with Tukey’s multiple comparisons test, significance values are denoted by: * p < 0.05; **, ## and $$ p < 0.001. **(B)** EM micrographs of paclitaxel−stabilized MTs alone and incubated with Tau or pTau. Scale bar = 100 nm for all images. **(C)** Representative total protein SDS-PAGE illustrating less pTau protein in pellet fraction compared to Tau (solid red arrows). sup. = supernatant, pel. = pellet.

Our results confirm the negative impact of tyrosine phosphorylation on the interaction of Tau with MTs, likely mediated by the large shift towards the negative electrostatic potential as well as structural rearrangements of MT-binding region through the phosphorylation of C-terminal tyrosine residues Y310 and Y394, with a possible contribution from other phosphorylated tyrosine residues requiring further investigation.

### Phosphorylation on tyrosine residues reduces Tau affinity for and binding of lipid vesicles

We have previously shown that both Tau and K18 interact with brain phosphatidylserine (BPS) vesicles resulting in vesicle disruption and the formation of highly stable oligomeric protein/phospholipid complexes (52). Therefore, we sought to assess whether phosphorylation on tyrosine residues influenced Tau interaction with membranous vesicles and the formation of such Tau-phospholipid complexes.

To address this question, we incubated Tau and K18 and their respective tyrosine-phosphorylated forms with BPS vesicles at a molar Tau to phospholipid ratio of 1 to 20 and performed sedimentation assay. We found that Tau and K18 incubated with BPS vesicles co−sedimented with vesicles to a larger extent compared to their phosphorylated counterparts. This is demonstrated by the reduction of phosphorylated Tau and K18 in the pellet fractions (Fig. 8, A, red box), signifying reduced Tau affinity and binding to lipids.

**Figure 8.**
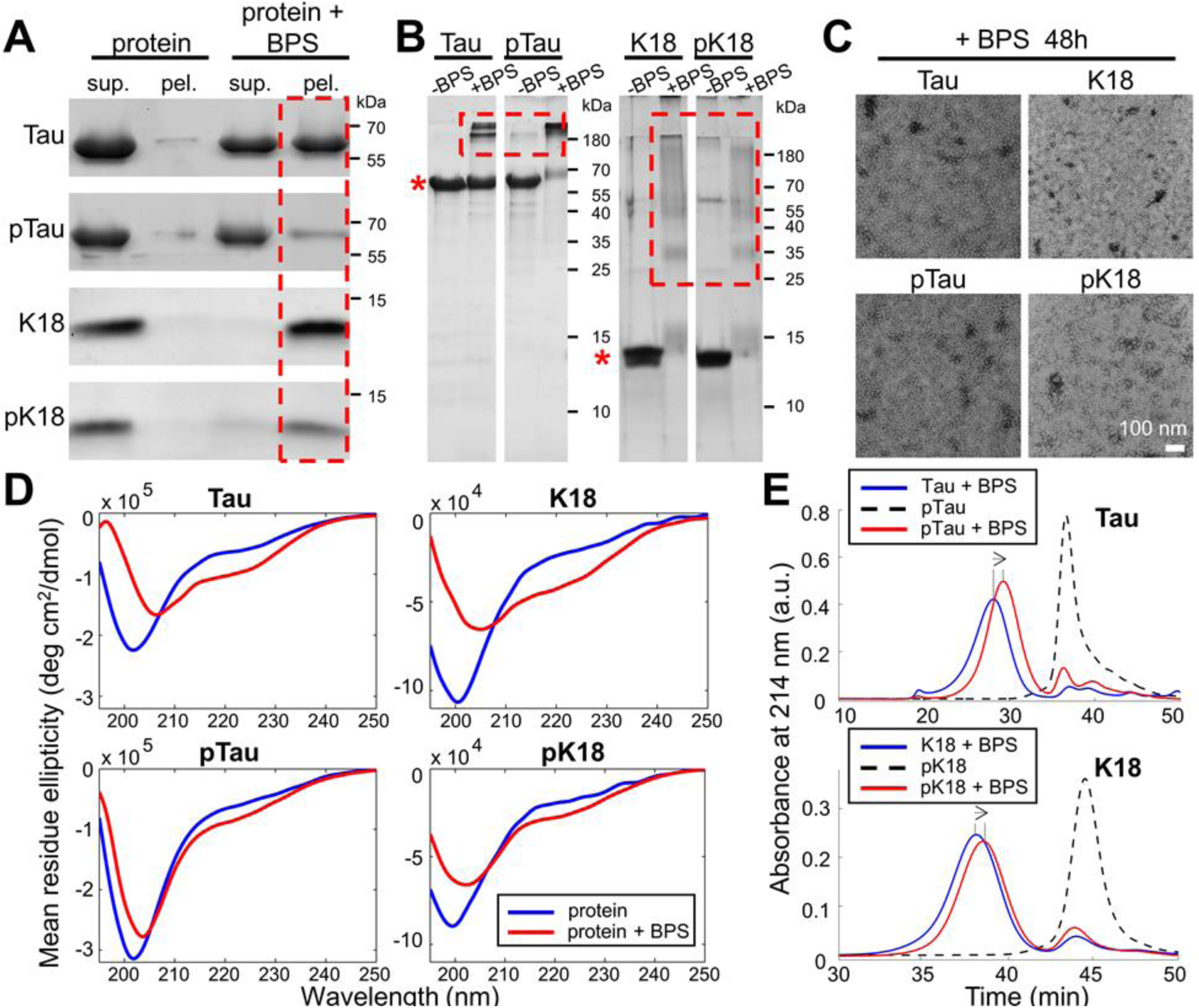
Phosphorylation on tyrosine residues reduces Tau affinity for and binding of lipid vesicles. **(A)** Co−sedimentation assay of BPS vesicles with 10 μM Tau, K18, pTau and pK18 at a molar ratio 1:20 (protein:phospholipid). **(B)** Native PAGE analysis of protein/phospholipid complexes formed by Tau, pTau (left) and K18, pK18 (right) after 24 h of incubation with BPS vesicles at a molar protein:phospholipid ratio of 1:20. The monomer bands are indicated by * and the presence of higher molecular weight bands indicated by red dashed rectangles. **(C)** EM micrographs showing that both WT and phosphorylated proteins were able to form phospholipid/protein complexes. Scale bar = 100 nm for all images. **(D)** CD spectra of proteins incubated for 2h alone (blue) or in presence of BPS vesicles at a molar protein:phospholipid ratio of 1:20 (red). **(E)** Assessment of the extent of Tau/K18-pTau/pK18-phospholipids complex formation using size−exclusion chromatography under the same conditions used in **(A)**. Dotted lines demarcate the elution peaks, arrows show the direction of the peak shift.

Next, to confirm the formation of oligomeric Tau- and K18-phospholipid complexes, Native gel analysis was performed, and showed the formation of oligomeric species by both non-phosphorylated and phosphorylated Tau and K18 (Fig. 8, B, red boxes). These findings were further supported by CD spectroscopy (Fig. 8, D), and EM, where oligomeric complexes were detected for all proteins without discernible morphological differences (Fig. 8, C), indicating no major differences in the absolute capacity of pTau and pK18 to form oligomeric structures compared to their non-phosphorylated counterparts.

Next, to verify the formation of the Tau/K18 phospholipid-oligomer complexes by the non-phosphorylated and phosphorylated Tau and K18, size exclusion chromatography (SEC) was performed. pTau and pK18 complexes peaks (Fig. 8, E, red) systematically eluted later compared to their non-phosphorylated counterparts (Fig. 8, E, blue; dashed lines indicate peaks of the elution profiles, arrows indicate the direction of elution profile shift) (Fig. 8, E, dashed). These data are well in line with biochemical data (Fig. 8, A, B) and suggest that Tau-lipid complex formation is affected by tyrosine phosphorylation of Tau and K18, resulting in oligomeric Tau- and K18-phospholipid complexes of smaller size, either through the reduction of the number of protein and/or phospholipid molecules per complex, through complex compaction. The differences in the pattern of the SDS-resistant oligomers formed by Tau and pTau oligomer suggests that they exhibit distinct biophysical properties and/or stability.

Tau and K18 were reported to undergo the structural transitions upon binding to lipid membranes (28,53-55). Therefore, to investigate the effects of tyrosine-phosphorylated Tau-phospholipids interactions, we monitored changes in Tau and K18 secondary structures by CD spectroscopy. Interestingly, the conformational transitions towards more structured forms of the proteins in the presence of BPS vesicles were markedly more pronounced for non-phosphorylated Tau and K18 (Fig. 8, D).

Our results demonstrate that phosphorylated Tau and K18 had reduced lipid vesicle binding affinity and formed oligomers that showed less pronounced structural conformation transitions and SEC elution profiles that were shifted towards smaller species compared to non-phosphorylated Tau and K18 in the presence of BPS vesicles. However, the capacity to associate with lipids was still preserved in the phosphorylated variants. Interestingly, the reduction of monomeric pTau protein band observed in BPS sample was concomitant with an increase of oligomeric bands of slightly different profile compared to those observed for non-phosphorylated Tau (Fig. 8, B). This observation combined with the differences in the SEC and CD profiles of phosphorylated Tau and K18 compared to the unmodified proteins suggest that the oligomers formed by the phosphorylated proteins exhibit distinct conformational and size properties.

### Tyrosine 310 phosphorylation attenuates Tau affinity for and binding of lipid vesicles

Next, we sought to determine the underlying structural basis for the reduced lipid-binding affinity of tyrosine-phosphorylated Tau. Previously, we showed that the core of the highly-stable oligomeric Tau-phospholipid complexes was comprised of both the PHF6* and PHF6 hexapeptide motifs, the latter in a β−sheet conformation (52). Since the Y310 residue is located in the PHF6 hexapeptide, part of the R3 repeat peptide (Fig. 1), which plays an important role in mediating the interaction between Tau and membranes (52,56), we set out to assess the effect of R3 Y310 phosphorylation on the lipid-mediated formation of fibrils by R3 and its interaction with membranes *in vitro*.

First, we incubated R3 and pR3 peptides with BPS vesicles at a molar peptide to phospholipid ratio of 1 to 1 and monitored fibril formation using EM (Fig. 9, A). We observed a very rapid formation of large fibrils by R3, whereas the pR3 peptide exhibited a significant delay in fibril formation (Fig. 9, A, 1h). Furthermore, whereas R3 formed long fibrils at a later time point, pR3 formed fibrils relatively shorter than R3 (Fig. 9, A, 24h, insets).

**Figure 9.**
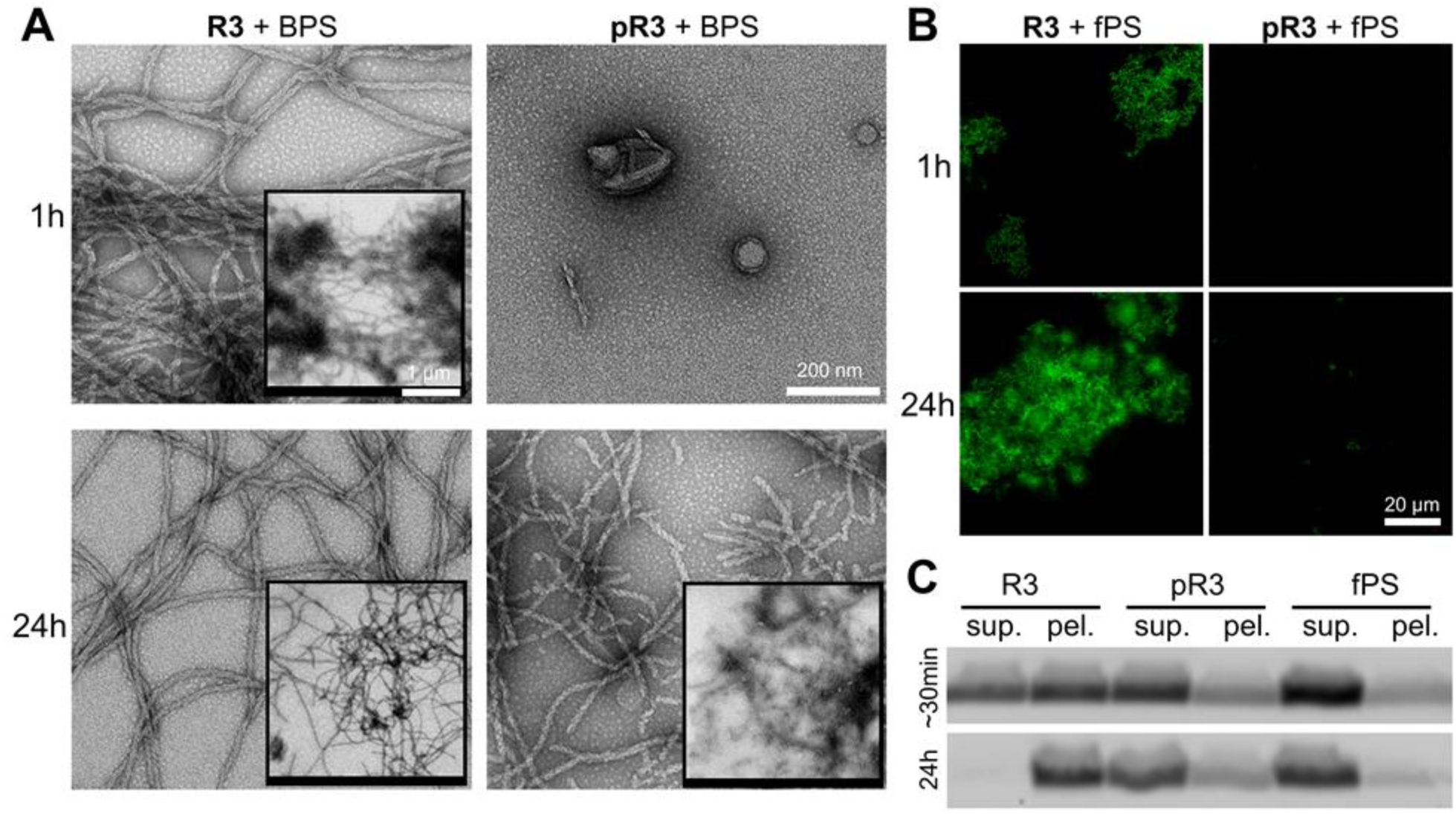
Tyrosine 310 phosphorylation attenuates R3 peptide affinity for and binding of lipid vesicles. **(A)** EM and **(B)** fluorescence microscopy images of 100 μM R3 and pR3 peptides incubated with BPS vesicles at a molar ratio of 1:1. For fluorescence microscopy, R3 or pR3 peptides were incubated with vesicles containing 1% fluorescent NBD−labelled phospholipids (fPS). Scale bar in **(A)** = 200 nm, in insets = 1 μm. Scale bar in **(B)** = 20 μm for all images. **(C)** SDS−PAGE gel analysis of the fluorescent lipid signal detection of the lipid vesicle flotation and peptide-lipid complex sedimentation assays. ∼30 min sample represents aliquots taken immediately after addition of peptides to fPS vesicles. Presence of faint signal in the pellet fraction of R3-fPS sample is likely due to fast association of peptide with lipids that occurred during the short period of time required to take and process the sample (estimated time ∼30min).

Our *in vitro* aggregation studies suggested that the phospholipids become incorporated within the fibrils during the fibrilization of R3. Therefore, we sought to assess the effects of Y310 phosphorylation influence R3 binding and incorporation of phospholipids into R3 fibrills. To this end, we co−incubated the R3 and pR3 peptides with vesicles containing 1% NBD−labelled fluorescent phospholipids (fPS). Under these conditions, we observed the formation of fluorescent fibrils in the case of R3, suggesting that R3 bound to and incorporated phospholipids within the fibril matrix. No fluorescent signal was observed in the case of the fibrils formed by pR3, suggesting that they were devoid of phospholipids (Fig. 9, B).

To further confirm the interaction of R3 and pR3 with the phospholipids, we performed a peptide-lipid sedimentation assay based on free lipid vesicle flotation. R3 and pR3 were incubated for 24 h with BPS vesicles containing 1% NBD−labelled fluorescent phospholipids, subsequently centrifuged, and run on SDS−PAGE and imaged with a fluorescent scanner, that detects fluorescent lipid signal (Fig. 9, C). In this assay, vesicles alone floated and remained in the supernatant (Fig. 9, C, fPS, sup.) and did not pellet to a high extent. However, upon association with the R3 peptide, a fluorescent signal could be observed in the pellet fractions. The signal present in the pellet fraction of R3-fPS at ∼30 min was likely due to the very aggregation and fibrillization of R3 that occurred during the short period of time required to aliquot and process the sample. We observed that phospholipids in the R3 sample were solely present in the pellet fraction after 24 h incubation, but that only a small fraction of the phospholipids in the pR3 sample were found in the pellet (Fig. 9, C). These results further confirm our EM and fluorescence microscopy data and indicate that the capacity of R3 to form fibrils that contain phospholipids in the presence of BPS vesicles was significantly diminished by phosphorylation of residue Y310. These findings are in line with data obtained with the full-length Tau and K18 data (Fig. 8), and further suggest that lipid incorporation into the fibrillar matrix is mediated by the aggregation−prone PHF6 peptide.

## Discussion

Despite the presence of tyrosine residues in domains that regulate Tau interactions with microtubules and its aggregation propensity, the role of Tau tyrosine phosphorylation in regulating Tau functions, structure, aggregation and toxicity remain generally understudied. For example, tyrosine 310 residue is located in the PHF6 motif of Tau protein, which has been identified as forming β-sheet structure in the core of Tau fibrils and is necessary for Tau aggregation *in vitro*. Despite consistent findings implicating PHF6 in the initiation of Tau aggregation, stabilization of Tau fibrils and the importance of the interaction of Y310 and I308 for Tau filament formation (35), there are no reports in the literature examining the potential impact of phosphorylation at this residue on the physiological and/or pathogenic properties of biologically-relevant full-length Tau proteoforms. This is possibly due to the lack of well-characterised pY310 specific antibodies and until recently the lack of methods for site-specific phosphorylation of Tau (57-59).

We speculated that modifications, such as the addition of bulky negatively-charged phospho-group to Y310 and/or other tyrosine residues would significantly alter the biophysical, aggregation and MT binding properties of Tau. In addition, the stoichiometry and localization of Tau tyrosine phosphorylation may confer differential effects on either N-terminal domain (Y18, Y29), that is implicated in Tau secretion (*60*) and regulation of microtubule stabilization (*61*), or on mid– and C-terminal domains (Y197, Y310, Y394), that are implicated in the modulation of Tau aggregation, microtubule- and lipid-binding properties (for a recent review see (62)). Furthermore, recent studies using cryoelectron microscopy of Tau filaments from human brain confirmed the PHF6 motif as a core component of Tau filaments, with Y310 buried within the interior of Tau filament in Pick’s disease, as well as in PHFs and SFs in AD, and Tau filaments in CTE (63-65) (Fig. 10), supporting our initial rationale for initiating the studies outlined in this manuscript. Although the extrapolations of *in vitro* studies’ results to *in vivo* conditions have obvious limitations and must be applied cautiously, the phosphorylation of Y310 residue, which is buried inside the core of Tau filaments in some Tauopathies, may have implications for some forms of disease featuring specific Tau strains.

**Figure 10.**
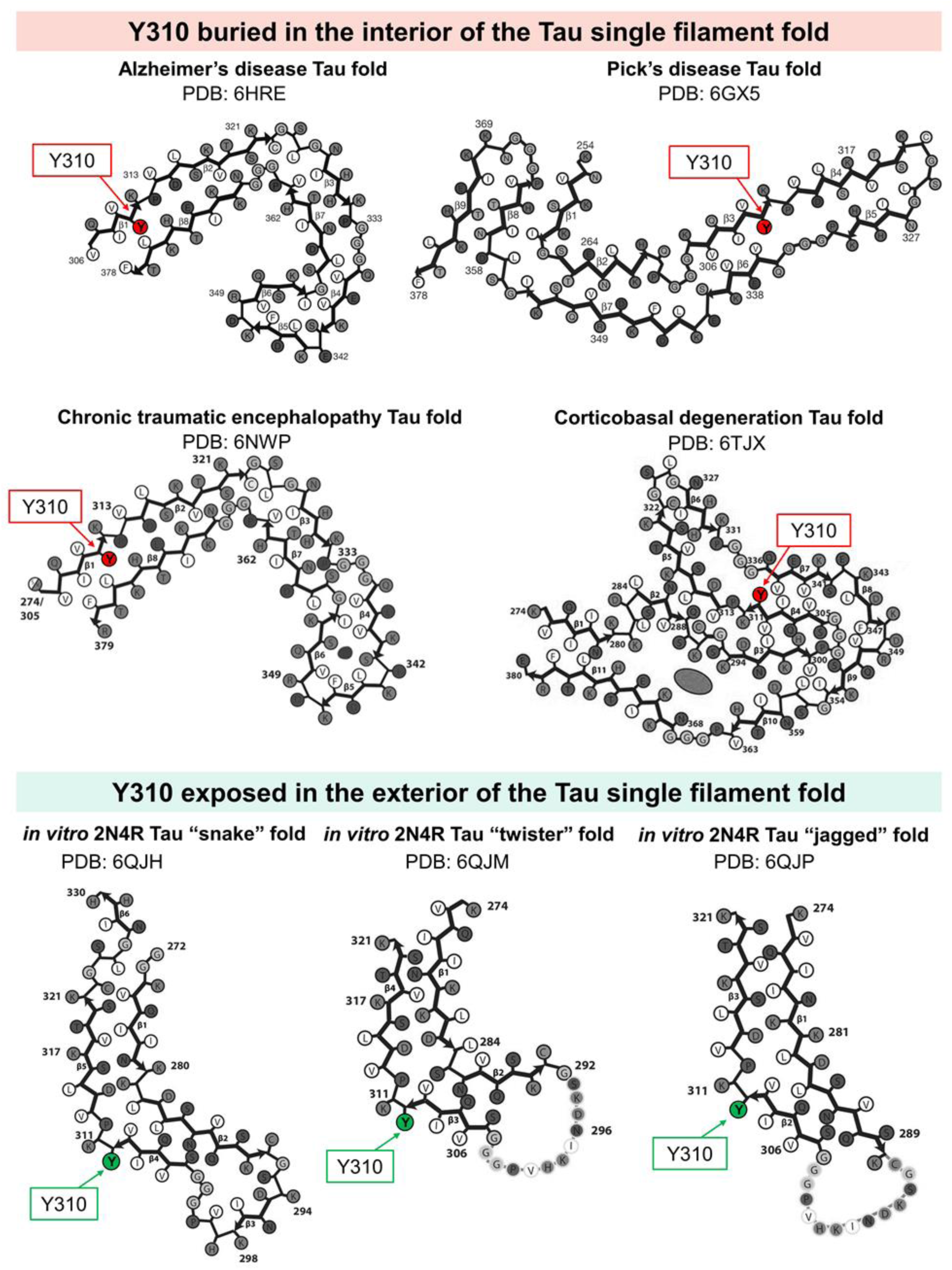
Schematic representations of Tau filament core fold from recently published Cryo-EM structures of brain-derived and *in vitro-*generated Tau filaments. Y310 residue is buried in the interior (red) of the Tau single filament fold in structures derived from AD, PiD, CTE and CBD-Tau, or exposed in the exterior (green) in *in vitro*-derived Tau structures. Adapted from (63-65,79,80).

Our results indicate that a) phosphorylation of Tau at multiple tyrosine residues inhibits Tau aggregation through cumulative effects of phosphorylation at multiple or all tyrosine residues, and that b) phosphorylation at residue Y310 is a key determinant of Tau aggregation *in vitro*. We show that Y310 phosphorylation appears to be sufficient to significantly delay Tau aggregation *in vitro* and reduces the propensity of the microtubule-binding region of Tau to aggregate through attenuation of the PHF6 hexapeptide propensity to adopt β-sheet rich structure in the context of the full-length Tau. Structural studies by NMR revealed that phosphorylation at Y310 induced local structural modifications in the PHF6 region of K18, thereby decreasing its propensity to adopt a β-sheet-containing secondary structure, contributing to the decreased aggregation propensity.

Y310 is the only tyrosine residue located in the MT-binding region of Tau, which highlights the potential role of Y310 in regulating Tau affinity for MTs. Our work here shows that phosphorylation on tyrosine residues affects full-length Tau affinity for MTs, in contrast to previous reports where Tau phosphorylated by Fyn (16), Arg (19) or c-Abl (18) *in vitro* was found to bind to MTs with an efficiency similar to that of the non-phosphorylated protein. These findings suggest that tyrosine phosphorylation of Tau on either Y18 (i.e. by Fyn) or Y394 (by c-Abl or Arg) did not significantly alter its affinity for MTs. Our data point toward residue Y310 as an important regulator of Tau/MTs interactions. One notable difference to previous reports is that our studies relied on the use of pTau proteins that are site-specifically phosphorylated at residues Y310 or Y394 (Figure 2, D). To the best of our knowledge, this is the first report demonstrating the role of tyrosine phosphorylation and Y310 in particular in regulating Tau MT-binding domain.

The fact that the MT-binding domain of Tau is also responsible for Tau-lipid interactions also points to the potential role of Y310 in regulating Tau affinity for lipid membranes. We have previously demonstrated that the PHF6 hexapeptide located within R3 interacts with negatively charged vesicles, resulting in the formation of β−strand—rich mixed phospholipid/peptide fibrils (52). In this work, we show that tyrosine phosphorylation, at least at residue Y310, reduced Tau binding to negatively-charged lipid membranes, which may have important implications for Tau functions at the cell plasma membrane. Our findings are especially relevant in the light of reports demonstrating that Tau tyrosine phosphorylation occurs in the human brain under both physiological and pathological conditions.

Although aberrant serine and threonine phosphorylation of Tau is well-established in the pathology of AD and other Tauopathies (66), the contributions of tyrosine phosphorylation to Tau pathology and Tau-mediated toxicity in these disorders remain poorly understood. Several lines of evidence suggest that Tau tyrosine phosphorylation is an early event in the development of AD (67). In 1991, Wood and Zinmeister used pan anti-phosphotyrosine antibodies to demonstrate the increase of non-specific tyrosine phosphorylation in AD patient’s brain tissue of NFT-bearing neuronal somatodendritic and dystrophic neurite compartments, as well as microglia-resembling cells within neuritic plaques, whereas predominantly neurite staining was observed in non-AD tissue (68). Similarly, Shapiro *et al*. reported increased non-specific phosphotyrosine immunoreactivity in the frontal cortex and hippocampus of AD patients (69). Since then, several reports based on antibody recognition or mass spectrometry have shown that Tau is phosphorylated on residues Y18, Y197 and Y394 (16,18,19,70) in AD and in foetal brain, whereas phosphorylation at positions Y18 (71) and Y394 (18) have been clearly shown to occur under physiological conditions.

Furthermore, in the P301L transgenic mouse model, Vega *et al*. showed that Tau phosphorylation at residues Y197 and Y394 correlated with the formation of Tau aggregates and occurred concurrently with serine and threonine phosphorylation (detected by PHF1 antibody at residues S396/S404 and by CP13 at residues S202/T205) known to be implicated in AD pathogenesis (70). Moreover, two tyrosine kinases, Fyn and c-Abl, which are known to phosphorylate Tau, have been shown to co-localize with Tau in NFTs (18,72). Interestingly, treatment with Aβ increased the activity of both kinases and led to a subsequent increase in Tau tyrosine phosphorylation and cell death, which could be prevented by treatment with tyrosine kinase inhibitors (73). In addition, Tau tyrosine phosphorylation may be involved in regulating several cellular activities of Tau linked to signalling events at the cell membrane, in lipid rafts and/or in the neuron growth cone (16,18). Overall, these findings strongly suggest that Tau tyrosine phosphorylation plays important roles in both the healthy state and in neurodegenerative disease conditions (74), and further studies investigating the specific role of Tau residue Y310 modifications are warranted.

In conclusion, our findings show that Tau phosphorylation on multiple tyrosine residues may have additive effects on inhibiting Tau fibril formation, and attenuating its MT- and lipid-binding affinity. In addition, our findings underscore the unique role of Y310 phosphorylation in regulating these properties of tau, which is consistent with its position within the aggregation-prone PHF6 hexapeptide, itself located in the MT-binding domain of the protein. We show that phosphorylation at Y310 is sufficient to significantly delay the fibrillization of full-length Tau and K18, and of the R3 repeat fragment. Our results strongly implicate tyrosine phosphorylation of Tau as a novel target for disease-modifying therapies of proteopathic disorders driven by Tau aggregation. Furthermore, our results underscore the critical importance of developing the new tools to assess the extent of Tau tyrosine phosphorylation in different Tauopathies and enable investigations of the role of tyrosine phosphorylation in regulating Tau function in health and disease.

## Experimental procedures

### DNA constructs and mutagenesis

The K18 fragments were synthesized by GeneArt Gene Synthesis (Life Technologies) and cloned into the pT7-7 for bacterial expression. Tau 2N4R in pCMV6-XL5 vectors was obtained from OriGene Technologies, Inc. and cloned in pET-15b for bacterial expression. Single site mutagenesis was performed in the pET-15b using the QuikChange Site-Directed Mutagenesis Kit (Stratagene), as per manufacturer’s instructions. pET21d SH2-CD c-Abl T231R and pCDFDuet-1 YopH for bacterial expression were kindly donated by Prof. Oliver Hantschel (ISREC Institute, EPFL).

### Peptides

The R3 (306-VQIVYKPVDLSKVTSKCGSLGNIHHK-331and the pY-R3 (306-VQIV(pY)KPVDLSKVTSKCGSLGNIHHK-331) corresponding to the third microtubule-binding repeat without/with phosphorylation at Y310 were purchased from CS Bio Co.

### Other compounds

Tubulin protein isolated from porcine brain (97% or >99 % pure), provided as a lyophilized powder was purchased from Cytoskeleton Inc. Paclitaxel from the pacific yew tree, *Taxus brevifolia* (purity of >99.5%), used inhibit microtubule depolymerization, provided as a lyophilized powder and resuspended at 2 mM anhydrous DMSO was purchased from Cytoskeleton Inc. (U.S.A). Guanosine triphosphate (GTP) provided as a lyophilized powder, resolubilized at 100 mM with ice-cold MilliQ water was purchased from Cytoskeleton Inc. (U.S.A). Microtubule (MT) associated protein fraction (MAPF) isolated from bovine brain, used as positive control for the MT binding assay, supplied as lyophilized powder was purchased from Cytoskeleton Inc. (U.S.A). Phosphatidylserine isolated from porcine brain (BPS) and fluorescently labeled phospholipid 1-oleoyl-2-{6-[(7-nitro-2-1,3-benzoxadiazol-4-yl)amino]hexanoylM-sn-glycero-3-phosphoserine (fPS) were purchased from Avanti Polar Lipids, Inc. Heparin sodium salt was from Applichem GmbH.

### Protein expression and purification (Tau and K18)

Tau isoforms 2N4R in pET-15b, K18 in pT7-7 were expressed in E. coli strain BL21. The purification protocol of Tau 2N4R Tau was adapted from (75). Cells were pelleted and broken by sonication in lysis buffer (3 M urea in 10 mM MES, pH 6.5, 1 mM DTT, 1 mM EDTA, 1 mM PMSF). After centrifugation at 150,000 g for 1 h at 4°C, 1% (w/v) of streptomycin sulfate was added to the supernatant, and the solution was stirred for 90 min at 4°C. After centrifugation at 27,000 g for 1 h at 4°C, the supernatant was dialyzed overnight at 4°C in ion exchange (IEX) buffer A (10 mM MES, pH 6.5, 20 mM NaCl, 1 mM DTT, 1 mM EDTA). The supernatant was filtered and loaded on a cation-exchange column (MonoS, GE Healthcare) and the protein was eluted using a salt gradient (increasing the NaCl concentration of IEX buffer A from 20 mM to 1 M NaCl over 20 column volumes). Fractions containing the proteins were dialyzed overnight against acetic buffer (5% acetic acid in water) and loaded on a reverse-phase HPLC C4 column (PROTO 300 C4 10 µm, Higgins Analytical; buffer A: 0.1% TFA in water, buffer B: 0.1% TFA in acetonitrile), and the protein was eluted using a gradient from 30 to 40% buffer B over 40 min (15 ml/min). K18 in pT7-7 was purified following a protocol adapted from (76). Briefly, cells were broken by sonication in IEX buffer B (10 mM HEPES, pH 6.5, 1 mM MgCl2, 20 mM NaCl, 1 mM DTT, 1 mM PMSF). After centrifugation at 40,000 g for 30 min, the supernatant was boiled until the solution became cloudy (∼5 min) and centrifuged again for 30 min. The supernatant was filtered and loaded on a cation-exchange column (MonoS, GE Healthcare), and the protein was eluted using a salt gradient (increasing the NaCl concentration of IEX buffer B from 20 mM to 1 M NaCl over 20 column volumes). Fractions containing the K18 fragment were immediately loaded on a reverse-phase HPLC C4 column (PROTO 300 C4 10 µm, Higgins Analytical; buffer A: 0.1% TFA in water, buffer B: 0.1% TFA in acetonitrile), and the protein was eluted using a gradient from 30 to 40% buffer B over 40 min (15 ml/min).

### SH2-CD c-Abl expression and purification

Recombinant SH2-CD c-Abl, T231R and YopH were co-overexpressed and SH2-CD c-Abl purified from E.coli as previously described (77). Briefly, cells were grown in 2 L of TB media, until the OD reached 1.0, after which the cultures were cooled down and induced overnight at 18°C, with 0.2 mM IPTG. The cells were then lysed by sonication in HisTag binding buffer HA (50 mM Tris pH 8, 500 mM NaCl, 5% Glycerol, 25 mM imidazole) and centrifuged twice at high speed (50 000 g, 20 min, 4°C), before being injected in 5 mL HisTag columns (histrap 5mL column, GE Healthcare, buffer HA: same as above, buffer HB: same as HA with 0.5 M imidazole). Selected fractions were desalted using two HiPrep 26/10 desalting columns (GE Healthcare) in series (Desalting buffer: 20 mM Tris pH 8, 50 mM NaCl, 5 % Glycerol, 1 mM DTT), combined and further purified by anion-exchange chromatography using a MonoQ 5/50 GL column (GE Healthcare, buffer A: 20 mM Tris, 5% Glycerol, 1 mM DTT, pH 8.0, buffer B: same as buffer A with 1 M NaCl). The final concentration of the c-Abl kinase was determined using the UV absorbance at 280 nm (M = 46797 g.mol-1 and ε280 = 80010 M-1.cm-1).

### Large scale preparation of phosphorylated K18, Tau and Y-> F mutants by c-Abl

The large-scale phosphorylation of 10 mg K18, Tau and Tau Y-> F mutants were performed for 4 h in 50 mM Tris, 5 mM MgCl2, 1 mM DTT, 20 mM Na3VO4 (phosphatase inhibitor) in the presence of 3 mM MgATP, pH,7.5 at 30°C. We used c-Abl kinase at a mass to mass ratio of 1:20 (kinase:Tau). The reaction mixture was followed by ESI-MS to verify the completion of the phosphorylation. Additional kinase and MgATP were added when needed. Phosphorylated K18, Tau and Y-> F mutants were purified by reverse-phase HPLC preparative C4 column (PROTO 300 C4 10 µm Higgins Analytical; buffer A: 0.1% TFA in water, buffer B: 0.1% TFA in acetonitrile) using a linear gradient of 30 to 40 % of B in 40 min for Tau or 20 to 35 % of B in 40 min for K18. The HPLC fractions were analyzed by LC-MS and once pure were pooled and lyophilized.

### Preparation and characterization of heparin-induced Tau and K18 fibrils

Fibrils of WT, mutants and phosphorylated Tau and K18 were formed by incubating monomeric protein in 10 mM phosphate, pH 7.4, 50 mM NaF and 0.5 mM DTT with heparin sodium salt (Applichem GmbH,Activity 204.7 I.U./mg MW: 8000-25000 g/mol) at a molar heparin:protein ratio of 1:4 under constant orbital agitation (1000 rpm, Peqlab, Thriller) at 37°C for up to 72 hours.

### R3 and pY-R3 peptide aggregation

100 µM R3 or pY-R3 peptide were dissolved in 10 mM Tris, 50 mM NaF,0.5 mM DTT and pH-adjusted to 7.4. The solution was then vortexed, centrifuged for 20 min at 14000 rpm and the supernatant was incubated in absence or presence of 25 µM heparin sodium salt (Applichem GmbH) at 37 °C, with constant orbital agitation (1000 rpm, Peqlab, Thriller).

### Vesicle preparations

Vesicles were prepared using the extrusion method. Briefly, individual phospholipids or phospholipid mixtures in chloroform were dried using an argon stream to form a thin film on the wall of a glass vial. Potentially remaining chloroform was removed by placing the vial under vacuum overnight. The phospholipids were resolubilized in 10 mM HEPES, pH 7.4, 100 mM NaCl and 2.5 mM DTT to the desired final concentration by sonication. The solution was then passed through an Avestin LiposoFast extruder (Avestin Inc.) (membrane pore size: 0.1 µm), and the size and homogeneity of the resulting vesicles were assessed by EM. For the preparation of fluorescently labeled vesicles, fluorescent and non-fluorescent phospholipids in chloroform were mixed at a molar ratio of 1:99 prior to the initial drying step.

### Preparation of mixed protein/phospholipid complexes

Protein/phospholipid complexes were prepared by incubating BPS vesicles with Tau or K18 at a molar protein:phospholipid ratio of 1:20 in 10 mM HEPES, pH 7.4, 100 mM NaCl, 2.5 mM DTT for 48 h at 37°C. The resulting protein/phospholipid complexes were separated from the remaining vesicles and soluble protein by SEC using a Superose 6 column (GE Healthcare).

### Preparation of mixed R3 and pY-R3 peptides/phospholipid fibrils

100 µM of R3 and pY-R3 peptides were incubated with BPS vesicles at a molar ratio of 1:1. For fluorescence microscopy, the peptides were incubated with vesicles containing 1% fluorescent NBD-phospholipids, imaged with confocal laser-scanning microscope (LSM 700, Zeiss) and analyzed using Zen software.

### Preparation of Microtubules

Tubulin was dissolved at a concentration of 5 mg/ml in ice-cold General tubulin buffer (80 mM PIPES, 0.5 mM EGTA, 2 mM MgCl2, pH 6.9) supplemented with 1 mM GTP and snap-frozen as 10 µl aliquots in liquid nitrogen and stored at -80°C. Microtubules were assembled by defrosting an aliquot of 10 µl tubulin stock in a RT water bath, immediately placed on ice and incubated for 20 min at 35°C in 100 µl pre-warmed (35°C) General tubulin buffer supplemented with 1 mM GTP and 5% glycerol. Then the solution was diluted with 100 µl General tubulin buffer supplemented with 1 µl of paclitaxel (Taxol©, stock at 2 mM), gently mixed and kept at RT. Microtubule morphology was systematically verified by electron microscopy.

### Microtubule binding assay

Pre-formed paclitaxel-stabilized MTs were incubated for 30 min at RT with Tau, pTau, MAPF or bovine serum albumin (BSA) (250 μg/ml protein and 100 μg/ml of tubulin) in General tubulin buffer supplemented with 1 mM GTP and 5% glycerol (total of 50 μl per sample). The samples were placed on top of a 100 μl Cushion buffer (i.e. General tubulin buffer containing 60 % glycerol) in 1.5 ml tubes. The tubes were centrifuged at 100’000 g for 40 min at RT. The top 50 μl of each tube was mixed with 10 μl of 5x Laemmli buffer (supernatant) and the pellet was resuspended in 50 μl 1x Laemmli buffer (pellet). The samples were run on SDS–PAGE gels. The relative amount of Tau, pTau, MAPF or BSA in the supernatant and in the pellet were estimated by measuring the band intensity using Fiji software (National Institute of Health) over 5 repeats, and represented as mean +/- SD.

### Monitoring of c-Abl Tau/K18 phosphorylation by mass spectrometry

100 or 200 μg of WT Tau or K18 were incubated with 5 or 10 μg of recombinant c-Abl in 50 mM Tris, 5 mM MgCl2, 1 mM DTT, 20 mM Na3VO4 (phosphatase inhibitor), 3 mM MgATP at a pH of 7.5. Reactions were performed for 4 h at 30°C without agitation. The reactions were followed using ESI–MS on a Thermo LTQ ion trap system.

### Tandem MS/MS of *in vitro* c-Abl phosphorylated recombinant Tau

Tau phosphorylated with C-Abl as described above were run on an SDS-PAGE and stained using Simply Blue Safe Stain (Life Technologies). The bands corresponding to Tau were excised, destained with 50% ethanol in 100 mM ammonium bicarbonate, pH 8.4, and dried. The proteins were digested in gel by covering gel pieces with a trypsin solution (12.5 ng/µl in 50 mM ammonium bicarbonate (pH 8.4), 10 mM calcium chloride) at 37°C overnight. The digested peptides were subjected to TiO2 phosphopeptide enrichment, concentrated to dryness, resuspended in acetonitrile and dried again. After repeating this step, the dried peptides were resuspended in 20 μl of 20% formic acid, and 2% acetonitrile, desalted on C18 OMIX tips (Agilent Technologies), and analyzed by a capillary LC–ESI–MS system (Thermo Scientific LTQ Orbitrap instrument). The collected MS/MS spectra were matched against human and scored using Mascot and Sequest algorithms. The data were then analyzed using Scaffold version 4.0 (Proteome Software, http://www.proteomesoftware.com/).

### Transmission electron microscopy

Samples (3.5 μl) were applied onto glow-discharged Formvar/carbon-coated 200-mesh copper grids (Electron Microscopy Sciences) for 1 min. The grids were blotted with filter paper, washed twice with ultrapure water and once with staining solution (uranyl formate 0.7% (w/V)), and then stained with uranyl formate for 30 sec. The grids were blotted and dried. Specimens were inspected with a Tecnai Spirit BioTWIN operated at 80 kV and equipped with an LaB6 filament and a 4K x 4K FEI Eagle CCD camera.

### Circular dichroism (CD) spectroscopy

CD spectra were recorded on a Jasco J-815 CD spectrometer operated at 20°C. To minimize buffer absorption, the buffer was modified to 10 mM phosphate buffer, pH 7.4, with 50 mM NaF and 0.5 mM DTT. CD spectra were acquired from 195 nm to 250 nm at a scan rate of 50 nm/min and in increments of 0.2 nm. For each sample, five spectra were averaged and smoothed using binomial approximation.

### Nuclear Magnetic Resonance (NMR) spectroscopy

^1^H-^15^N HSQC spectra were collected at 25 °C on a Bruker Avance 600 MHz spectrometer using samples containing 500 μM K18 or pK18 peptide in 10 mM Na_2_HPO_4_, 100 mM NaCl, 1 mM DTT, 10% D_2_O buffer at pH 7.4. Resonance assignments are based on the previously published assignments of the K18 spectrum (66).

### Sedimentation assay

Taking advantage of the fact that high-speed centrifugation sediments fibrils but not monomeric or soluble oligomeric species, the sedimentation assay makes it possible to assess the ratio of fibrillised-to-soluble protein. 20 µl of WT Tau, K18 or Y->F mutants aggregation reactions were pelleted by ultracentrifugation at 200’000 g of 2 h at 4°C. The supernatant was mixed with 2x Laemmli buffer and run on 15% SDS-PAGE. When stated, the amount of soluble material was quantified by measuring the band intensity using Fiji software (National Institute of Health) and averaged over at least three independent repeats.

### Thioflavin T fluorescence measurements

Thioflavin T (ThT) fluorescence reading (excitation wavelength of 450 nm, emission wavelength of 485 nm) was performed in triplicate with a ThT concentration of 60 μM and a peptide concentration of 60 μM or a protein concentration of 3 μM in 50 mM glycine, pH 8.5, using a Bucher Analyst AD plate reader without shaking.

### Vesicle binding assay

The co-sedimentation assay was used to assess initial binding of protein to vesicles. Tau or K18 WT, or tyrosine phosphorylated (10 µM) were mixed with BPS vesicles (200 µM) to a total volume of 20 µl and immediately centrifuged at 200,000 g for 2 h at 4°C. At this speed, the vesicles pellet, and protein bound to them co-sediment with the vesicles. Both the supernatant, containing unbound protein, and the resuspended pellet, containing vesicles with bound protein, were run on 15% SDS-PAGE gels.

## Acknowledgements

This work was supported by funding from the Swiss Federal Institute of Technology Lausanne (H.A.L.) and partially by AC Immune. Support was also provided by NIH grant R37AG019391 to D.E. NMR was performed at the Weill Cornell NMR core facility, supported by NIH grant S10OD016320.We would like to thank Dr. Prof Olivier Hantschel (EPFL) for providing us with the c-Abl kinase plasmid, Romain Hamelin and the proteomic core facility (EPFL) for the MS/MS data acquisition. We also would like to acknowledge the EPFL CIME facility providing access to EM. We thank Nathalie Jordan and Céline Vocat for their excellent technical support with cloning and protein expression/purification, respectively.

## FOOTNOTES

## The abbreviations used are

Abl: Abelson murine leukemia viral oncogene homolog 1
AD: Alzheimer’s disease
Arg: Abelson tyrosine-protein kinase 2
BPS: brain phosphatidylserine
CBD: corticobasal degeneration
CD: circular dichroism
CTE: chronic traumatic encephalopathy
EM: electron microscopy
fPS: fluorescent phospholipids
FTDP-17: frontotemporal dementia with parkinsonism linked to chromosome 17
GTP: guanosine triphosphate
HSCQS: heteronuclear single quantum coherence spectroscopy
LC/MS: liquid chromatography–mass spectrometry
MAPF: microtubule−associated protein fraction
MS/MS: tandem mass spectrometry
MT: microtubule
MTBD: microtubule-binding domain
MTBR: microtubule-binding region
NBD: nitrobenzoxadiazole
NFT: neurofibrillary tangle
NMR: nuclear magnetic resonance
PD: Parkinson’s disease
pS: phospho-serine
PSP: progressive supranuclear palsy
pT: phospho-threonines
PTM: post-translational modification
pY: phospho-tyrosine
RP−HPLC: reversed phase high-performance liquid chromatography
RT: room temperature
SD: standard deviation
SDS-PAGE: sodium dodecyl sulphate-polyacrylamide gel electrophoresis
SP: spectral count
ssi: secondary shift index
Syk: Spleen tyrosine kinase
ThT: thioflavin T
TTBK1: Tau tubulin kinase 1
UPLC: ultra performance liquid chromatography
WT: wild-type

